# Neonatal immune challenge induces female-specific changes in social behavior and somatostatin cell number, independent of microglial inflammatory signaling

**DOI:** 10.1101/2020.04.30.068924

**Authors:** Caroline J. Smith, Marcy A. Kingsbury, Julia E. Dziabis, Richa Hanamsagar, Karen E. Malacon, Jessica N. Tran, Haley A. Norris, Mary Gulino, Staci D. Bilbo

## Abstract

Decreases in social behavior are a hallmark aspect of acute “sickness behavior” in response to infection. However, immune insults that occur during the perinatal period may have long-lasting consequences for adult social behavior by impacting the developmental organization of underlying neural circuits. Microglia, the resident immune cells of the central nervous system, are sensitive to immune stimulation and play a critical role in the developmental sculpting of neural circuits, making them likely mediators of this process. Here, we investigated the impact of a postnatal day (PND) 4 lipopolysaccharide (LPS) challenge on social behavior in adult mice. Somewhat surprisingly, neonatal LPS treatment decreased sociability in adult female, but not male mice. LPS-treated females also displayed reduced social interaction and social memory in a social discrimination task as compared to saline-treated females. Somatostatin (SST) interneurons within the anterior cingulate cortex (ACC) have recently been suggested to modulate a variety of social behaviors. Interestingly, the female-specific changes in social behavior observed here were accompanied by an increase in SST interneuron number in the ACC. Finally, these changes in social behavior and SST cell number do not appear to depend on microglial inflammatory signaling, because microglia-specific genetic knock-down of myeloid differentiation response protein 88 (MyD88; the removal of which prevents LPS from increasing proinflammatory cytokines such as TNFα and IL-1β) did not prevent these LPS-induced changes. This study provides novel evidence for enduring effects of neonatal immune activation on social behavior and SST interneurons in females, independent of microglial inflammatory signaling.

## 1. Introduction

Social interactions are essential to survival, health, and well-being across species. However, in the event that individuals are exposed to pathogens such as viruses and bacteria, it is critical that social contact be acutely limited. Indeed, decreased social motivation is a hallmark adaptive component of “sickness behavior” in response to immune activation (Dantzer & Kelley, 2007). Conversely, prolonged or developmental immune challenges could lead to maladaptive and lasting deficits in social behavior. It is increasingly recognized that the immune system has the capacity to modulate neural circuits both during states of immune activation, as well as under normal, healthy circumstances (Bluthe et al., 1994; Konsman et al., 2008; Schafer et al., 2012; Filiano et al., 2016). This is especially true during development, as microglia, the resident immune cells of the brain, play a critical role in sculpting neural circuits, including those that support social behavior (Nelson et al., 2017; Kopec et al., 2018; VanRyzin et al., 2019). Importantly, many disorders that are characterized by social dysfunction, such as autism spectrum disorder (ASD), often have an immune component to their etiology (McDougle et al., 2015; Gottfried et al., 2015). Furthermore, changes in both microglial morphology, gene expression, and maturity have been identified in individuals with social dysfunction (Hanamsagar et al., 2017; Gandal et al., 2018; Morgan et al., 2014; Werling et al., 2016).

Within the neural networks supporting social behavior, the anterior cingulate cortex (ACC) is a subdivision of the pre-frontal cortex (PFC) that represents a critical node, across species (Bicks et al., 2015; Guo et al., 2020; Apps et al., 2016). Importantly, both the structure and functional connectivity of the ACC are altered in individuals with ASD (Postema et al., 2019; Guo et al., 2020; Laidi et al., 2019; Zhou et al., 2016). Somatostatin (SST) interneurons are important for maintaining and regulating excitatory/inhibitory balance and this cell population within the ACC has recently been implicated in the modulation of a variety of social behaviors including social interaction (Perez et al., 2019; Sun et al., 2020; van Heukelum et al., 2019), social affective discrimination (Scheggia et al., 2020), socio-sexual behavior (Nakajima et al., 2014), and social fear learning (Xu et al., 2019) in animal models. Thus, we hypothesized that perinatal immune stimulation with lipopolysaccharide (LPS) would induce social behavior impairments and decrease SST interneuron number in the ACC in adulthood in mice.

One critical factor when considering the relationship between immune activation and social behavior is biological sex. A large body of evidence demonstrates that the response to an immune challenge, as well as its enduring consequences for the brain and behavior, differs between the sexes (for review see Klein & Flanagan, 2016; Hanamsagar & Bilbo, 2016). These sex differences in neuroimmune function are present during development, as evidenced by differences in microglial gene expression, morphology, maturity, and responses to immune challenge (see comprehensive reviews by Bordt et al., 2019; VanRyzin et al., 2019) and may lead to sex differences in disease susceptibility. For example, males are at much higher risk of immune-related neurodevelopmental disorders such as ASD (Baio et al., 2018) while females are far more likely than males to develop autoimmune disorders such as lupus erythematosus, myasthenia gravis, and thyroid diseases including Graves disease and Hashimotos thyroiditis (Klein & Flanagan, 2016), as well as anxiety and depression (Hodes & Epperson, 2019). While several studies have investigated the impact of neonatal LPS challenge on behavioral outcomes and/or microglia (Peng et al., 2019; Bukhari et al., 2018; MacRae et al., 2015; Williamson et al., 2011; Schwarz & Bilbo, 2011; Rico et al., 2010), few of these studies have included both males and females in their analysis, especially in the context of social behavior (Taylor et al., 2012; Kentner et al., 2018; Carlezon et al., 2019). Given that females have traditionally been underrepresented in such studies (Beery & Zucker, 2011; Klein & Flanagan, 2016) it is critical that we gain a better understanding of the long-term consequences of neonatal inflammation in females.

We predicted that neonatal immune challenge would act via microglia-mediated immune processes to induce behavioral deficits, potentially in a sex-specific manner. To manipulate microglial toll-like receptor (TLR) proinflammatory signaling, and thus, the microglial response to neonatal LPS, we utilized a novel transgenic mouse recently developed and characterized in our lab. In this mouse model, myeloid differentiation response protein 88 (MyD88) gene expression is knocked down only in CX3CR1-positive tissue resident macrophages, i.e. microglia within the brain (Rivera et al., 2019). MyD88 is an adaptor protein downstream of most TLRs, with the exception of TLR3 (Akira & Takeda, 2004). *In vitro*, MyD88-depleted microglia isolated from these transgenic mice fail to produce the proinflammatory cytokines tumor necrosis factor alpha (TNFα) and interleukin-1 beta (IL-1β) in response to LPS (Rivera et al., 2019). Therefore, our second hypothesis was that any effects of LPS on the brain and behavior would be prevented by removal of microglial-MyD88 signaling.

Surprisingly, we report here that a neonatal LPS challenge reduced social behavior in adult female, but not male mice. This was accompanied by an increase in SST cell number in the ACC. Furthermore, although MyD88 knockdown significantly decreased TNFα and IL-1β mRNA in microglia following an LPS challenge, this genetic manipulation had no significant effects on either social behavior or SST cell number in females. Taken together, these results demonstrate that neonatal endotoxin exposure has enduring consequences for social behavior in females, independent of microglial-MyD88 signaling.

## Methods

### 2.1 Animals

MyD88-floxed mice were purchased from Jackson Laboratories (Bar Harbor, ME; Stock # 00888). Cx3Cr1-CreBT (MW126GSat) mice were generated and provided by L. Kus (GENSAT BAC Transgenic Project, Rockefeller University, NY) and backcrossed over 12 generations on a C57Bl/6N Charles River background. Wild-type (WT) C57Bl/6J mice were purchased from Jackson Laboratories (Bar Harbor, ME; Stock # 000664). All methods for breeding and genotyping transgenic mice were conducted according to Rivera et al., 2019. Cx3Cr1-CreBT mice were crossed with MyD88-floxed mice over 3 generations until all offspring were fully MyD88-floxed (F/F) and either Cre negative (Cre ^0/0^: no modification to microglial MyD88) or Cre positive (Cre ^tg/0^: removal of microglial MyD88). Genotyping of transgenic animals was conducted using polymerase chain reaction (PCR) on tail-snip DNA. Primer sequences used for PCR can be found in Table 1. All animals were group housed in standard mouse cages under standard laboratory conditions (12-hour light/dark cycle, 23°C, 60% humidity) with same-sex littermates. All experiments were conducted in accordance with the NIH *Guide to the Care and Use of Laboratory Animals* and approved by the Massachusetts General Hospital Institutional Animal Care and Use Committee (IACUC).

**Table 1.**
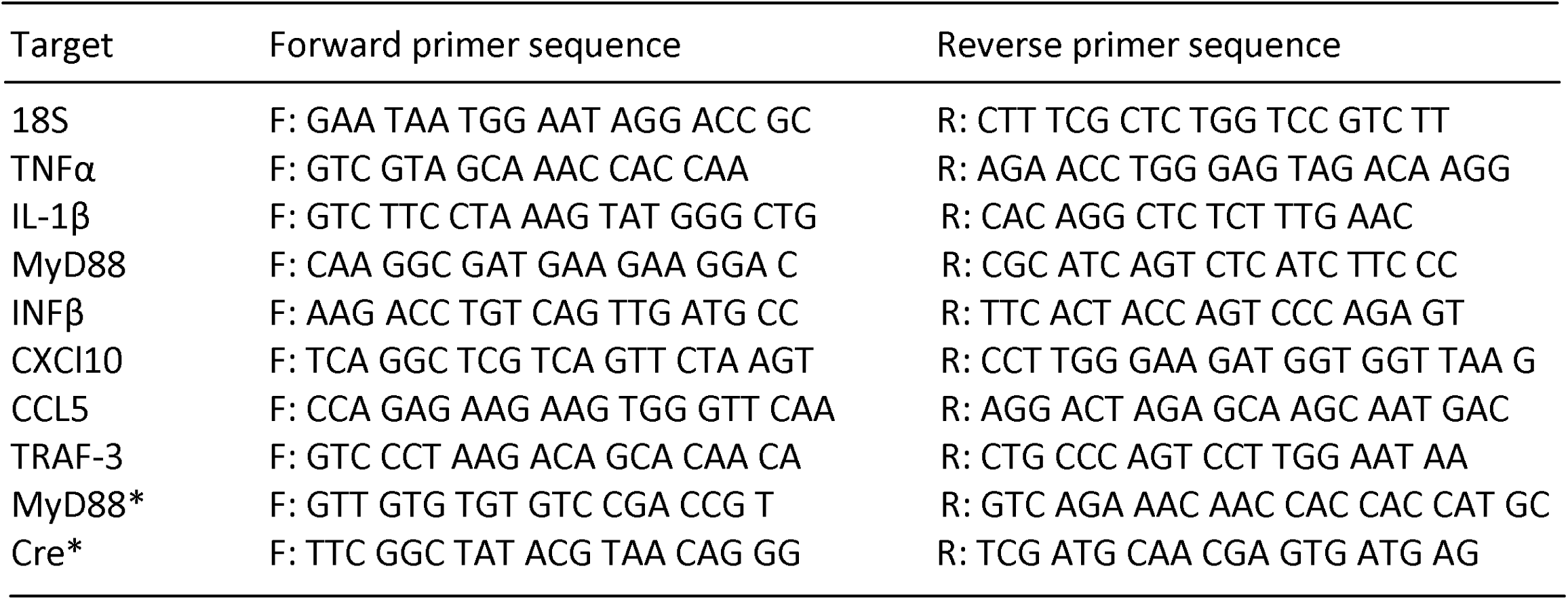
Primers used for qPCR and PCR (genotyping). * denotes primers used for genotyping.

### 2.2 Neonatal LPS challenge

330 µg/kg LPS (from Escherichia coli serotype 0111:B4, Sigma-Aldrich, St. Louis, MO, USA) dissolved in 0.9% sterile saline or 0.9% sterile saline control (SAL) were administered subcutaneously on postnatal day (PND) 4 in all experiments. This dose was chosen based on previous work demonstrating its sufficiency to induce changes in microglial morphology and gene expression (Hanamsagar et al., 2017) and PND 4 was chosen because we have previously found that immune challenge at this timepoint induces long lasting changes in microglial function (Williamson & Bilbo, 2014; Schwarz & Bilbo, 2011). All animals within a litter received the same treatment to prevent any indirect exposure to the opposite drug treatment.

### 2.3 Microglial Isolations

In order to verify the efficacy of our transgenic model to reduce proinflammatory gene expression in microglia following an immune challenge, microglial isolations were conducted according to Hanamsagar et al. 2017. Briefly, microglia were isolated from bilateral hippocampi on PND 4, 2 hours after LPS administration. Hippocampi were dissected from whole brains on ice using sterile forceps and minced with a razor blade. Homogenate was then placed in Hank’s Buffered Salt Solution (HBSS; Thermofisher Scientific, NY, USA) containing collagenase A (Roche, Catalog #: 11088793001, Indianapolis, IN, USA; 1.5mg/ml) and DNAse 1 (Roche, Catalog #: 10104159001, Indianapolis, IN, USA; 0.4 mg/ml) and incubated in a water bath at 37°C. Mechanical digestion of tissue was performed by sequentially passing samples through successively smaller glass Pasteur pipettes. Once a single cell suspension was obtained, samples were filtered, rinsed in HBSS, and centrifuged at 2400 rpm for 10 min at 4°C. Next, samples were incubated with CD11b antibody conjugated magnetic beads (Miltenyi Biotec, Catalog #: 130-093-634 San Diego, CA) for 15 min and then passed through a magnetic bead column (Quadro MACS Separator and LS columns, Miltenyi Biotec, Catalog #: 130-042-401, San Diego, CA) to separate CD11b+ cells (microglia) from CD11b-cells. Both CD11b+ and CD11b-cell populations were washed in 1X PBS and stored at −80°C until RNA extraction. This is a well-established method for isolating microglia from brain tissue (see Bordt et al., 2020, STAR protocols).

### 2.4 RNA extraction

Samples were homogenized in Trizol® (Thermo-fisher scientific, NY) and then vortexed for 10 min at 2000 rpm. After 15 min resting at room temperature (RT), chloroform was added (1:5 with Trizol) and samples were vortexed for 2 min at 2000 rpm. Samples were next allowed to sit at RT for 3 min and then centrifuged (15 min at 11,800 rpm; 4°C). This resulted in a gradient from which the top, clear, aqueous phase was separated and placed in to a fresh tube. Isopropanol was added to the new tube to precipitate RNA (1:1 with aqueous phase) and samples were again vortexed, allowed to set at RT for 10 min, and then centrifuged (15 min at 11,800 rpm; 4°C). Pellets obtained after this step were rinsed twice in ice-cold 75% Ethanol and then resuspended in 8 µl of nuclease-free water. RNA was frozen at −80°C until cDNA synthesis and qPCR. A NanoDrop Spectrophotometer (Thermo Scientific, Wilmington, DE) was used to determine RNA quantity and purity. RNA was considered pure enough for further use based on 260/280 (RNA:protein; range: 1.8-2.0) and 260/230 (RNA: Ethanol; range 1.6-2.0) ratios.

### 2.5 cDNA synthesis & qPCR

cDNA was synthesized using the QuantiTect Reverse Transcription Kit (Quiagen, Hilden, Germany). Briefly, 200ng of RNA/12 µl of nuclease-free water was pre-treated with gDNase at 42°C for 2 min to remove genomic DNA contamination. Next, master mix containing both primer-mix and reverse transcriptase was added to each sample and all samples were heated to 42°C for 30 min and then 95°C for 3 min (to inactivate the reaction) in the thermocycler. Quantitative real-time PCR (qPCR) was conducted using QuantiTect SYBR Green PCR Kit (Quiagen, Catalog #:204057, Hilden, Germany). For gene expression analyses, we selected the MyD88 gene and two genes downstream of MyD88, TNFα and IL-1β, to verify knockdown of MyD88-dependent signaling that leads to the transcription of pro-inflammatory cytokines via NF-_Κ_B activation (Table 1; see also McCarthy et al., 2017). To examine MyD88-independent signaling that is mediated by the TRIF-dependent pathway, leading to the activation of interferon (INF) and interferon inducible genes, we selected 4 genes downstream of TRIF that included TRAF3, INFβ, CCL5 and CXCL10 (Table 1; see also MaCarthy et al., 2017). qPCR primers were designed in the lab and ordered from Integrated DNA technologies (Coralville, IA). All primer sequences can be found in Table 1. PCR product was monitored using a Mastercycler ep *realplex* (Eppendorf, Hauppauge, NY). Male and female samples were run on separate plates, and thus, cannot be directly compared. I8S was used as a house-keeping gene and relative gene expression was calculated using the 2^-^ΔΔ^CT^ method, relative to 18S and the lowest sample on the plate (Williamson et al., 2011; Livak & Schmittgen, 2001).

### 2.6 Behavioral Assays

Behavioral assays were conducted in adulthood, in a separate cohort of animals from those used for PCR analyses, at approximately 10 weeks of age. For all assays, testing took place in the latter half of the light phase. Animals were moved to the behavioral testing room 1 hour prior to behavioral testing and were habituated to the testing room for 1 afternoon prior to the first day of testing. Estrus cycle phase was assessed in all females using cell characterizations from vaginal smears. All behavior was video recorded and scored using Solomon Coder (https://solomon.andraspeter.com) by an observer who was blind to the experimental conditions of the animals.

#### 2.6.1 Social Preference

To assay the preference of experimental animals to investigate a social vs a non-social stimulus, a 3-chambered social preference task was used according to Smith et al., 2015. In this task, the testing apparatus consists of a 3-chambered box with openings to allow for passage from one chamber to the next. Separate 3-chambered boxes were used for males and females, and sexes were tested on separate days to prevent interference from opposite-sex odors. Stimuli (either a novel sex- and age-matched conspecific or a novel rubber duck) were confined within smaller containers composed of Plexiglass rods in each of the opposite side chambers. Experimental animals were placed into the middle chamber and their movement and investigation of each of the stimuli was scored over the course of 5 min. All animals were habituated to the testing apparatus without the social and object stimuli for 5 min on the day before testing. The testing apparatus was cleaned with a disinfectant between each test. Scored elements were: time spent in each chamber, time spent investigating each stimulus (i.e. direct sniffing or nose-poking between the bars of the smaller stimulus containers), and time spent in the middle empty chamber. In order to quantify social preference, ‘sociability’ was calculated as a proportion of investigation time spent investigating the social stimulus ((social investigation time / (social + object investigation time)) x 100).

#### 2.6.2 Social Discrimination

To test social memory, a social discrimination test was used according to Lukas et al., (2013) and Engelmann et al (1995) with some modifications. This test takes advantage of the fact that mice prefer to investigate novel conspecifics over ones that are familiar. In a clean, novel cage, female experimental animals were first exposed to a novel female juvenile conspecific for 5 min. Following this exposure, experimental animals were singly-housed in a separate clean cage for an interval of 30 min. This interval was chosen to fall within the time window at which adult mice have previously been shown to retain a social memory (Lukas et al., 2013). After this interval, they were again exposed to the first juvenile stimulus animal (now familiar) along with a second novel animal for 5 min. During this 5 min period behaviors scored were anogenital investigation of either stimulus animal, whole body investigation of either stimulus animal, rearing, digging, and autogrooming. A social discrimination score was calculated for both whole body investigation ((novel investigation time / (novel + familiar investigation time)) x 100) and anogenital investigation ((novel anogenital investigation time / (novel + familiar anogenital investigation time)) x 100). In order to assess social discrimination at the initial reintroduction, social discrimination scores (whole body and anogenital) were also separately measured during the first 1 min of the test.

### 2.7 Tissue Collection & Sectioning

In a final cohort of behaviorally naïve female animals treated neonatally with LPS or saline, brain tissue was collected at PND 60 for immunohistochemical analysis. All animals were euthanized using C0_2_ inhalation. Transcardial perfusion with 0.9% saline was used to clear blood from tissue and brains were removed and post-fixed in 4% paraformaldehyde for 48 hrs at 4°C followed by 30% sucrose for 48 hrs at 4°C. Next, brains were rapidly frozen in 2-methylbutane on dry ice and stored at −80°C until sectioning. Brains were cut on a Leica cryostat into 40 µm coronal sections and stored in cryoprotectant until immunohistochemical processing.

### 2.8 Somatostatin Immunohistochemistry & Cell Quantification

Immunohistochemistry was used to label somatostatin-positive interneurons in the ACC. Briefly, sections were removed from cryoprotectant and rinsed 4 × 5 min in 1X PBS. Next, prior to primary antibody labeling, tissue was pretreated with 0.3% H_2_0_2_ for 10 min, 10 mM Sodium Citrate (pH 9.0) for 30 min at 75°C, and then 10% goat serum (blocking) for 1 hr, with 1X PBS rinses between each step. Tissue was exposed to rabbit anti-somatostatin (Somatostatin-14, 1:1000, Lot: A17908, Peninsula, San Carlos, CA) overnight at 4°C, followed by secondary antibody for 1 hr the next day (AlexaFluor goat anti-rabbit 488, 1:500, Thermofisher Scientific). Sections were mounted onto gelatin-subbed slides and coverslipped with vectashield anti-fade mounting medium (Vector Labs, Burlingame, CA).

SST cell number was quantified in the ACC at 3 “Levels” designated based on Bregma distances (Paxinos & Watson, 5^th^ Edition; See Figure 5 for visual representation). ‘Level 1’ corresponds with 1.13 mm anterior to Bregma, ‘Level 2’ corresponds with 0.73 mm anterior to Bregma, and ‘Level 3’ corresponds with 0.03 mm anterior to Bregma. Cumulatively this covers the majority of the rostro-caudal extent of the ACC. Z-stack images (19 Z planes at a 1.68 µm interval per field) were captured on a Zeiss AxioImager.M2 Microscope at 10x, using a 2×3 tile with 10% overlap to cover the entire region of interest. A maximum intensity projection was generated for each tiled image in which all SST neurons were counted in the entire ACC at all three rostro-caudal levels and expressed as number of cells per defined area. Cell counts were performed by an experimenter blind to the genotype and treatment condition of the experimental animals.

### 2.9 Statistics

All statistics and data visualization were conducted using GraphPad Prism Version 8. For qPCR analyses, a 3-way ANOVA (genotype x treatment x cell type) was conducted for each gene of interest in each sex separately. For sociability and social discrimination testing, 2-way ANOVA’s (genotype x treatment) were used to analyze behavior. In the sociability test, male and female data were analyzed separately because animals were tested at separate times. 2-way ANOVA (genotype x treatment) was used to analyze somatostatin cell number. Bonferroni posthoc tests were conducted for multiple comparisons. All group N’s are reported in the figure legends. Statistical significance was set at p<0.05.

## 2. Results

### 3.1 Removal of microglial-MyD88 signaling blunts MyD88-dependent, but not MyD88-independent, proinflammatory gene expression in response to PND4 LPS challenge, specifically in microglia

In both males and females, as expected, MyD88 mRNA was significantly higher in CD11b+ cells (microglia) as compared to CD11b-cells (Males: F _(1, 22)_ = 78.12, p<0.0001; Females: F_(1, 19)_ = 169.2, p<0.0001) and significantly lower in Cre ^tg/0^ (MyD88 knockdown) animals than in Cre ^0/0^ animals (Males: F_(1, 27)_ = 9.479, p<0.01; Females: F_(1, 30)_ = 28.22, p<0.0001). Furthermore, MyD88 mRNA did not differ between Cre ^0/0^ and Cre ^tg/0^ animals in CD11b-cells (Males: F_(1, 22)_ = 11.30, p<0.01; Females: F_(1, 19)_ = 28.33, p<0.0001), consistent with the microglial specificity of this genetic manipulation (see Table 2 for complete statistics; Fig. 1A&B).

**Table 2.** 3-way ANOVA analysis of qPCR results. All main and interaction effects for each gene of interest are represented for both males and females. Bolded statistics indicated significant effects.

| Gene | Cell Type | Genotype | Treatment | Genotype x Cell Type | Genotype x Treatment | Cell Type x Treatment | Genotype x Cell Type x Treatment |
| --- | --- | --- | --- | --- | --- | --- | --- |
| **Males** |  |  |  |  |  |  |  |
| *MyD88-dependent genes* |  |  |  |  |  |  |  |
| MyD88 | $F_{(1, 22)} = \mathbf{78.12, p<0.0001}$ | $F_{(1, 27)} = \mathbf{9.479, p<0.01}$ | $F_{(1, 27)} = \mathbf{12.85, p<0.01}$ | $F_{(1, 22)} = \mathbf{11.30, p<0.01}$ | $F_{(1, 27)} = 0.031, p=0.86$ | $F_{(1, 22)} = \mathbf{8.841, p<0.01}$ | $F_{(1, 22)} = 0.197, p=0.66$ |
| IL-1β | $F_{(1, 26)} = \mathbf{75.81, p<0.0001}$ | $F_{(1, 26)} = \mathbf{10.45, p<0.01}$ | $F_{(1, 26)} = \mathbf{40.20, p<0.0001}$ | $F_{(1, 26)} = \mathbf{10.20, p<0.01}$ | $F_{(1, 26)} = 3.841, p=0.055$ | $F_{(1, 26)} = \mathbf{38.97, p<0.0001}$ | $F_{(1, 26)} = 3.744, p=0.059$ |
| TNFα | $F_{(1, 25)} = \mathbf{61.24, p<0.0001}$ | $F_{(1, 27)} = \mathbf{11.17, p<0.01}$ | $F_{(1, 27)} = \mathbf{25.97, p<0.0001}$ | $F_{(1, 25)} = \mathbf{11.25, p<0.01}$ | $F_{(1, 27)} = 3.312, p=0.080$ | $F_{(1, 25)} = \mathbf{24.34, p<0.0001}$ | $F_{(1, 25)} = 23.382, p=0.078$ |
| *MyD88-independent genes* |  |  |  |  |  |  |  |
| TRAF3 | $F_{(1, 26)} = \mathbf{4.462, p<0.05}$ | $F_{(1, 27)} = 0.002, p=0.961$ | $F_{(1, 27)} = \mathbf{5.362, p<0.05}$ | $F_{(1, 26)} = 3.342, p=0.79$ | $F_{(1, 27)} = 0.372, p=0.547$ | $F_{(1, 26)} = 3.910, p=0.059$ | $F_{(1, 26)} = 0.862, p=0.771$ |
| CCL5 | $F_{(1, 26)} = \mathbf{12.99, p<0.01}$ | $F_{(1, 27)} = 0.018, p=0.890$ | $F_{(1, 27)} = \mathbf{10.75, p<0.01}$ | $F_{(1, 26)} = 0.037, p=0.848$ | $F_{(1, 27)} = 0.009, p=0.927$ | $F_{(1, 26)} = \mathbf{10.99, p<0.01}$ | $F_{(1, 26)} = 0.022, p=0.883$ |
| CXCL10 | $F_{(1, 26)} = \mathbf{35.51, p<0.0001}$ | $F_{(1, 27)} = 0.216, p=0.646$ | $F_{(1, 27)} = \mathbf{50.51, p<0.0001}$ | $F_{(1, 26)} = 0.005, p=0.947$ | $F_{(1, 27)} = 0.325, p=0.573$ | $F_{(1, 26)} = \mathbf{31.22, p<0.0001}$ | $F_{(1, 26)} = 0.045, p=0.833$ |
| INFβ | $F_{(1, 26)} = \mathbf{16.79, p<0.001}$ | $F_{(1, 27)} = 0.969, p=0.334$ | $F_{(1, 27)} = \mathbf{8.224, p<0.01}$ | $F_{(1, 26)} = 0.832, p=0.370$ | $F_{(1, 27)} = 1.509, p=0.230$ | $F_{(1, 26)} = \mathbf{8.222, p<0.01}$ | $F_{(1, 26)} = 1.369, p=0.253$ |
| **Females** |  |  |  |  |  |  |  |
| *MyD88-dependent genes* |  |  |  |  |  |  |  |
| MyD88 | $F_{(1, 19)} = \mathbf{169.2, p<0.0001}$ | $F_{(1, 30)} = \mathbf{28.22, p<0.0001}$ | $F_{(1, 30)} = \mathbf{17.19, p<0.001}$ | $F_{(1, 19)} = \mathbf{28.33, p<0.0001}$ | $F_{(1, 30)} = \mathbf{8.546, p<0.01}$ | $F_{(1, 19)} = \mathbf{4.666, p<0.05}$ | $F_{(1, 19)} = \mathbf{7.601, p<0.05}$ |
| IL-1β | $F_{(1, 16)} = \mathbf{174.4, p<0.0001}$ | $F_{(1, 33)} = \mathbf{8.392, p<0.01}$ | $F_{(1, 33)} = \mathbf{94.06, p<0.001}$ | $F_{(1, 16)} = \mathbf{8.396, p<0.05}$ | $F_{(1, 33)} = \mathbf{9.243, p<0.01}$ | $F_{(1, 16)} = \mathbf{95.35, p<0.0001}$ | $F_{(1, 16)} = \mathbf{9.315, p<0.01}$ |
| TNFα | $F_{(1, 19)} = \mathbf{55.41, p<0.0001}$ | $F_{(1, 30)} = \mathbf{10.61, p<0.01}$ | $F_{(1, 30)} = \mathbf{34.54, p<0.001}$ | $F_{(1, 19)} = \mathbf{9.927, p<0.01}$ | $F_{(1, 30)} = \mathbf{12.59, p<0.01}$ | $F_{(1, 19)} = \mathbf{32.47, p<0.0001}$ | $F_{(1, 19)} = \mathbf{11.83, p<0.01}$ |
| *MyD88-independent genes* |  |  |  |  |  |  |  |
| TRAF3 | $F_{(1, 19)} = \mathbf{5.379, p<0.05}$ | $F_{(1, 30)} = 0.000, p=0.996$ | $F_{(1, 30)} = 0.154, p=0.697$ | $F_{(1, 19)} = 0.029, p=0.866$ | $F_{(1, 30)} = 0.144, p=0.707$ | $F_{(1, 19)} = 0.142, p=0.711$ | $F_{(1, 19)} = 1.466, p=0.241$ |
| CCL5 | $F_{(1, 18)} = \mathbf{42.46, p<0.0001}$ | $F_{(1, 30)} = 0.305, p=0.585$ | $F_{(1, 30)} = \mathbf{25.39, p<0.0001}$ | $F_{(1, 18)} = 0.174, p=0.682$ | $F_{(1, 30)} = 4.051, p=0.053$ | $F_{(1, 18)} = \mathbf{22.87, p<0.001}$ | $F_{(1, 18)} = 3.447, p=0.080$ |
| CXCL10 | $F_{(1, 19)} = \mathbf{8.262, p<0.01}$ | $F_{(1, 30)} = 0.466, p=0.500$ | $F_{(1, 30)} = \mathbf{34.24, p<0.0001}$ | $F_{(1, 19)} = 0.137, p=0.716$ | $F_{(1, 30)} = 0.569, p=0.457$ | $F_{(1, 19)} = \mathbf{5.741, p<0.05}$ | $F_{(1, 19)} = 0.216, p=0.647$ |
| INFβ | $F_{(1, 19)} = \mathbf{8.383, p<0.01}$ | $F_{(1, 30)} = 0.001, p=0.972$ | $F_{(1, 30)} = 2.147, p=0.153$ | $F_{(1, 19)} = 0.034, p=0.857$ | $F_{(1, 30)} = 2.560, p=0.120$ | $F_{(1, 19)} = 1.627, p=0.218$ | $F_{(1, 19)} = 1.992, p=0.174$ |

**Figure 1.**
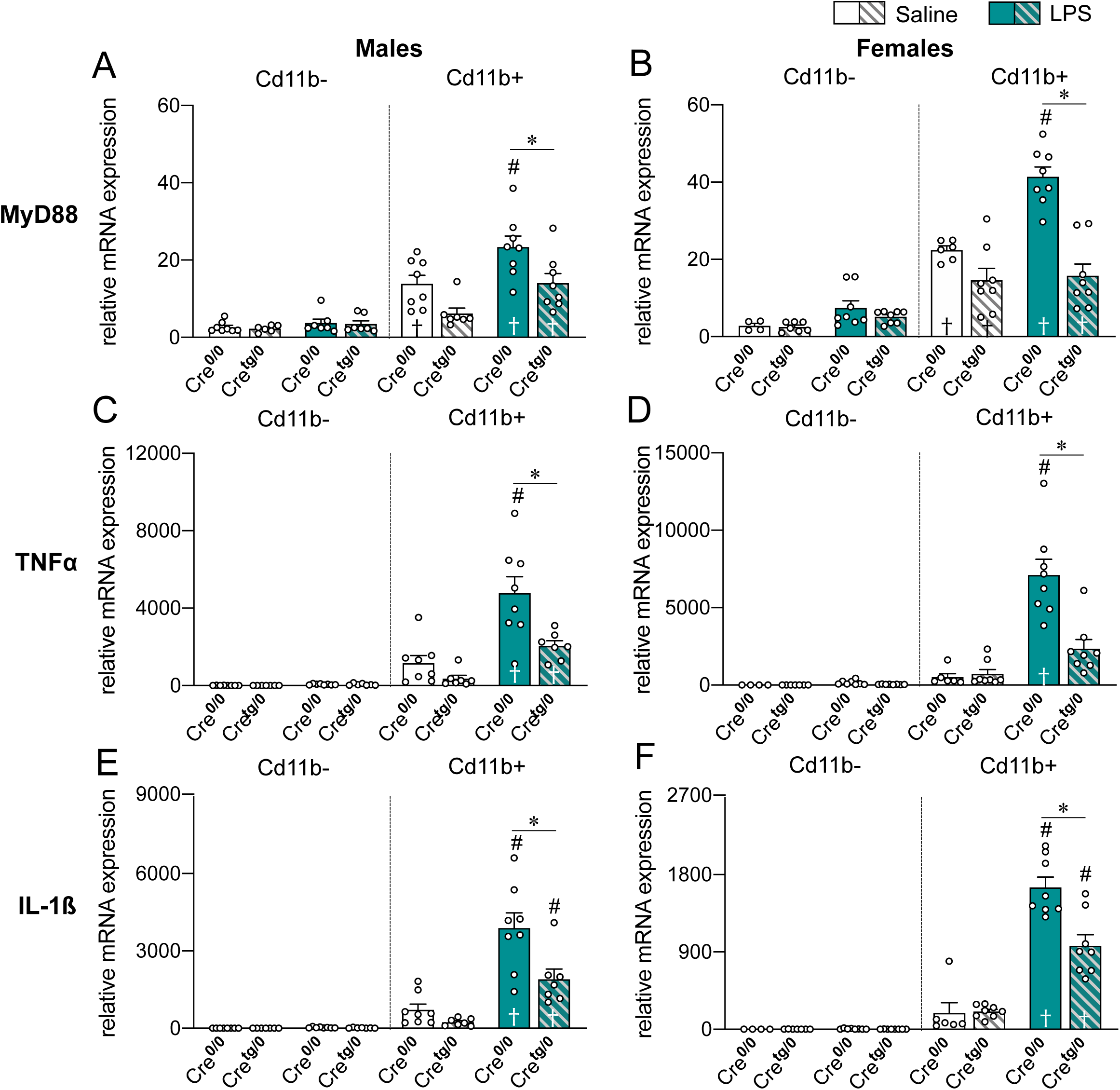
Removal of microglial-MyD88 signaling blunts MyD88-dependent proinflammatory gene expression in response to P4 LPS challenge, specifically in microglia. In males, LPS-treatment increases the gene expression in Cre^0/0^animals, but not Cre^tg/0^ animals, of MyD88 (**A**), TNFα (**C**), and IL-1β (**E**) in CD11b+ microglia. In females, LPS-treatment also fails to increase MyD88 mRNA (**B**), and TNFα mRNA (**D**) in Cre^tg/0^ animals, and induces significantly less IL-1β (**F**) in microglia. Results of 3-way ANOVA (cell type x treatment x genotype), *Bonferroni *posthoc* effects of genotype, ^#^Bonferroni posthoc effects of treatment (relative to saline counterpart), †Bonferroni posthoc effects of cell type (relative to CD11b-counterpart), p<0.05.

**Figure 2.**
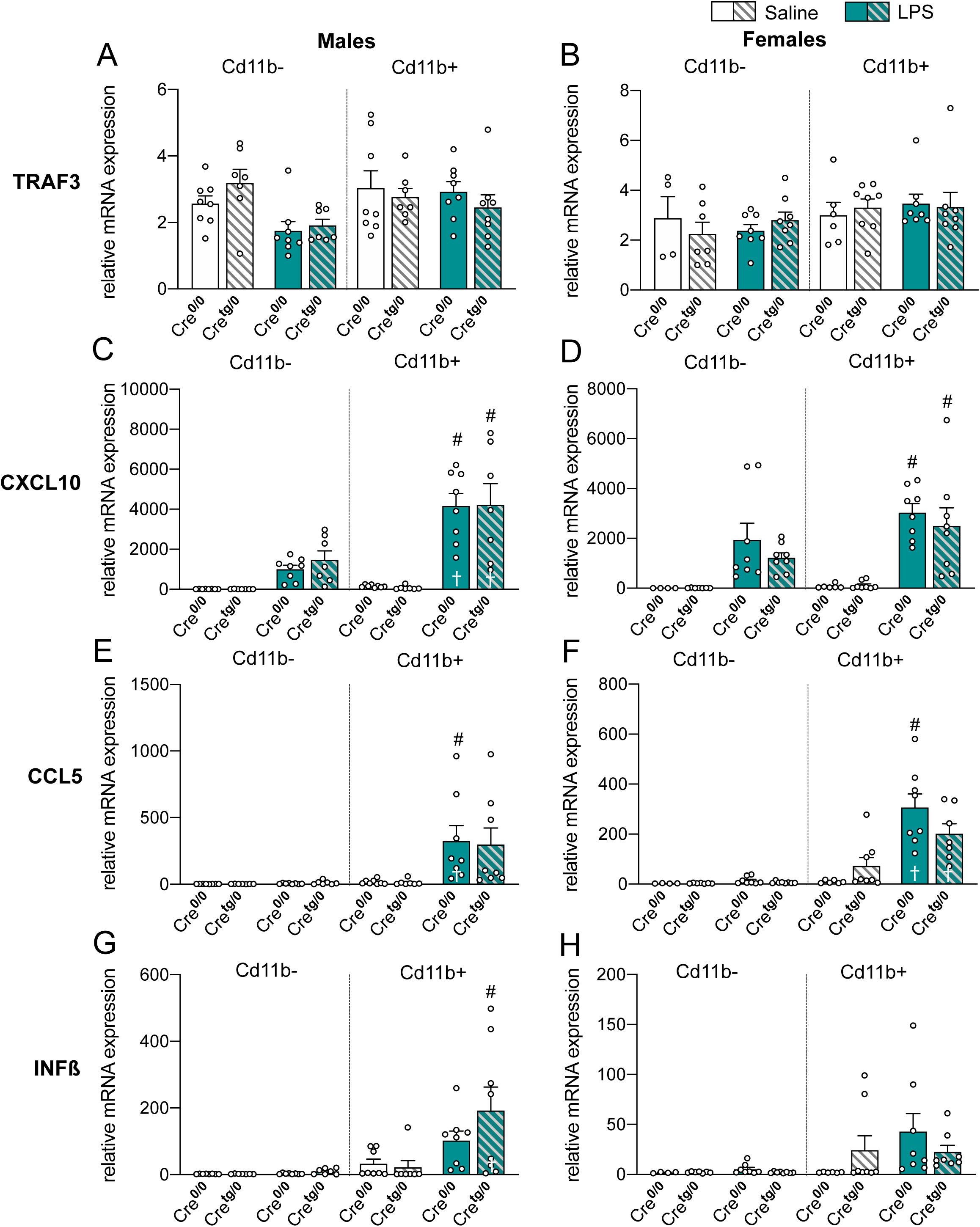
Removal of microglial-MyD88 signaling has no effect on MyD88-independent gene expression in response to PND4 LPS challenge, specifically in microglia. No genotype effects were observed in TRAF3 (**A**,**B**), CXCL10 (**C**,**D**), CCL5 (**E**,**F**) or INFβ (**G**,**H**) in either sex. Results of 3-way ANOVA (cell type x treatment x genotype), *Bonferroni *posthoc* effects of genotype, ^#^Bonferroni posthoc effects of treatment (relative to saline counterpart), †Bonferroni posthoc effects of cell type (relative to CD11b-counterpart), p<0.05.

Significant main effects of cell type, genotype, and treatment were observed for MyD88-dependent proinflammatory cytokines downstream of TLR activation in both sexes (See Table 2 for complete main and interaction effects; Fig1C-F). Specifically, in both males and females, TNFα mRNA expression was higher in microglia as compared to CD11b-cells (Males: F_(1, 25)_ = 61.24, p<0.0001; Females: F_(1, 19)_ = 55.41, p<0.0001). LPS treatment significantly increased microglial TNFα mRNA (Males: F_(1, 27)_ = 25.97, p<0.0001; Females: F_(1, 30)_ = 34.54, p<0.001). Posthoc testing revealed that this LPS-induced increase was blunted in Cre ^tg/0^ animals (Males: p<0.001; Females: p<0.0001) and did not differ from saline treatment (Males: p=0.09; Females: p=0.45). Similarly, IL-1β mRNA expression was higher in microglia as compared to CD11b-cells (Males: F _(1, 26)_ = 75.81, p<0.0001, Females: F_(1, 16)_ = 174.4, p<0.0001). Posthoc testing showed that this LPS-induced increase was also blunted in Cre ^tg/0^ animals (Males: p<0.001; Females: p<0.0001). These results demonstrate the successful knockdown of MyD88-dependent signaling within microglia in Cre ^tg/0^ animals.

In contrast, while mRNA for the MyD88-independent genes CXCL10, CCL5, INFβ, and TRAF3 was more highly expressed in microglia as compared to CD11b-cells, there were no significant main effects of genotype in either males or females (See Table 2 for complete main and interaction effects; Fig2A-F). LPS treatment increased CXCL10, CCL5, and INFβ mRNA in males (CXCL10: F_(1, 27)_ = 50.51, p<0.0001; CCL5: F_(1, 27)_ = 10.75, p<0.01; INFβ: F_(1, 27)_ = 8.224, p<0.01; TRAF3: F_(1, 27)_ = 5.362, p<0.05) and CXCL10 and CCL5 mRNA in females (CXCL10: F_(1, 30)_ = 34.24, p<0.0001; CCL5: F_(1, 30)_ = 25.39, p<0.0001), but these increases were not prevented in Cre ^tg/0^ animals. These results demonstrate that MyD88 deletion did not alter microglial gene expression in the TRIF-dependent interferon pathway, nor did it augment the activity of this pathway (e.g. in a compensatory fashion).

### 3.2 PND4 LPS challenge decreases social preference in adulthood, in females but not males, regardless of microglial-MyD88 signaling

Given the literature suggesting male-biased sensitivity to early-life immune challenges, we predicted that males would be more robustly impacted by LPS challenge than females, and that these behavioral effects would be prevented in Cre ^tg/0^ animals that have blunted proinflammatory signaling. Interestingly, however, neither LPS treatment nor genotype had significant effects on any measure of social preference (Fig. 3A) in males (see Table 3 for complete statistics; Fig. 3B-F). However, in females, neonatal LPS treatment significantly decreased sociability, regardless of genotype (F _(1, 31)_ = 4.533, p<0.05, Fig. 3C). LPS treatment also decreased time spent in the social chamber (F _(1, 31)_ = 7.27, p<0.05, Fig. 3E), and increased time spent in the object chamber (F _(1, 31)_ = 4.924, p<0.05, Fig. 3G). For social chamber time, there was a trend towards an increase in Cre ^tg/0^ animals as compared to Cre ^0/0^ animals (F _(1, 31)_ = 3.402, p=0.075). No differences were observed in total investigation time, or in middle chamber time (Table 3).

**Table 3.** 2-way ANOVA results of sociability testing in males and females. Bolded statistics indicate significant effects, * based on first 1 min. of the test; s : seconds.

| Behavior | Treatment | Genotype | Interaction |
| --- | --- | --- | --- |
| **Males:** |  |  |  |
| Social chamber time (s) | $F_{(1, 33)} = 1.966, p=0.170$ | $F_{(1, 33)} = 0.000, p=0.990$ | $F_{(1, 33)} = 0.285, p=0.597$ |
| Object chamber time (s) | $F_{(1, 33)} = 1.321, p=0.258$ | $F_{(1, 33)} = 0.004, p=0.948$ | $F_{(1, 33)} = 0.001, p=0.976$ |
| Middle chamber time (s) | $F_{(1, 33)} = 0.326, p=0.572$ | $F_{(1, 33)} = 0.024, p=0.877$ | $F_{(1, 33)} = 1.323, p=0.258$ |
| Sociability (%) | $F_{(1, 34)} = 0.000, p=0.986$ | $F_{(1, 34)} = 0.000, p=0.994$ | $F_{(1, 34)} = 0.716, p=0.406$ |
| Social investigation (s) | $F_{(1, 33)} = 0.442, p=0.511$ | $F_{(1, 33)} = 0.022, p=0.882$ | $F_{(1, 33)} = 0.073, p=0.789$ |
| Object investigation (s) | $F_{(1, 33)} = 2.691, p=0.110$ | $F_{(1, 33)} = 0.775, p=0.395$ | $F_{(1, 33)} = 0.590, p=0.448$ |
| Total investigation (s) | $F_{(1, 33)} = 1.515, p=0.227$ | $F_{(1, 33)} = 0.033, p=0.857$ | $F_{(1, 33)} = 0.286, p=0.596$ |
| **Females:** |  |  |  |
| Social chamber time (s) | **$F_{(1, 31)} = 7.27, p<0.05$** | $F_{(1, 31)} = 3.402, p=0.075$ | $F_{(1, 31)} = 0.171, p=0.682$ |
| Object chamber time (s) | **$F_{(1, 31)} = 4.924, p<0.05$** | $F_{(1, 31)} = 2.639, p=0.114$ | $F_{(1, 31)} = 0.017, p=0.899$ |
| Middle chamber time (s) | $F_{(1, 31)} = 0.870, p=0.358$ | $F_{(1, 31)} = 0.178, p=0.676$ | $F_{(1, 31)} = 0.318, p=0.577$ |
| Sociability (%) | **$F_{(1, 31)} = 4.533, p<0.05$** | $F_{(1, 31)} = 1.659, p=0.207$ | $F_{(1, 31)} = 0.319, p=0.576$ |
| Social investigation (s) | $F_{(1, 31)} = 3.383, p=0.076$ | $F_{(1, 31)} = 0.935, p=0.341$ | $F_{(1, 31)} = 0.299, p=0.588$ |
| Object investigation (s) | $F_{(1, 31)} = 1.828, p=0.186$ | $F_{(1, 31)} = 1.086, p=0.305$ | $F_{(1, 31)} = 0.177, p=0.677$ |
| Total investigation (s) | $F_{(1, 31)} = 0.341, p=0.564$ | $F_{(1, 31)} = 1.001, p=0.973$ | $F_{(1, 31)} = 0.025, p=0.876$ |

**Figure 3.**
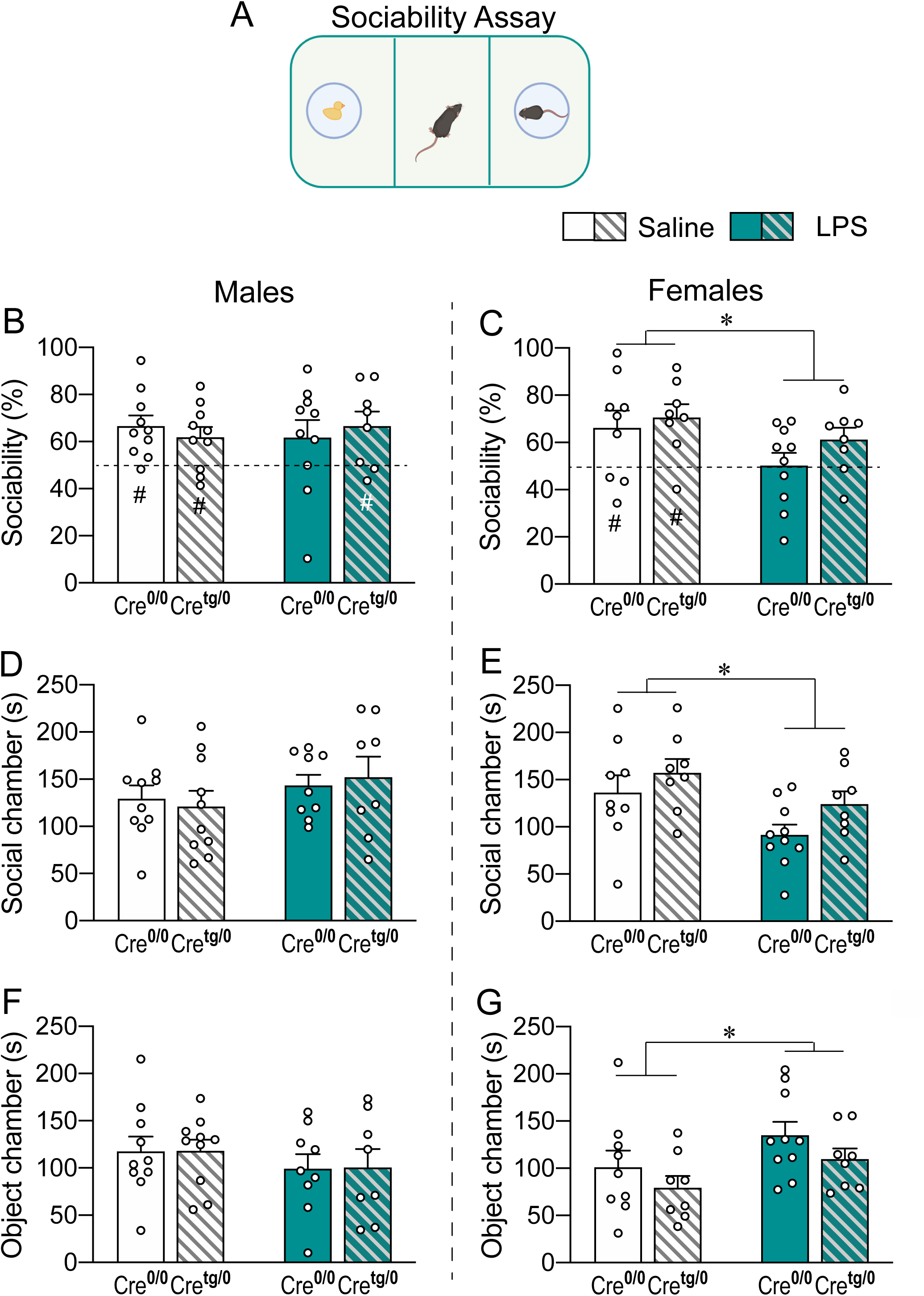
PND4 LPS challenge decreases social preference in adulthood, in females but not males, regardless of microglial-MyD88 signaling. **A**) Sociability Assay in which the subject is given the opportunity to interact with either a novel sex- and age-matched conspecific or a novel object in a 3-chambered apparatus for 5 min. Neither LPS treatment nor genotype had any significant effects on male sociability (**B**), social chamber time (**D**), or object chamber time (**F**). In contrast, neonatal LPS treatment significantly decreased sociability (**C**) and social chamber time (**E**) in females, while increasing object chamber time (**G**). *main effects of 2-way ANOVA (treatment x genotype), p<0.05. ^#^single sample t-test against 50% (chance), p<0.05.

### 3.3 PND4 LPS challenge reduces social discrimination in females, independent of microglial-MyD88 signaling

To further characterize the female-specific social deficits induced by neonatal LPS treatment, social discrimination was tested in females (see Table 4 for complete statistics). During the full 5 min of the social discrimination trial of the test (Trial 2, Fig 4A), LPS-treated females spent less time in total investigation (F _(1, 33)_ = 4.686, p<0.05, Fig. 4B), novel investigation (F _(1, 33)_ = 5.471, p<0.05, Fig. 4C), and anogenital investigation of the novel stimulus animal (F _(1, 33)_ = 4.321, p<0.05, Fig. 4D), as compared to saline-treated females, regardless of genotype. There were no significant differences in familiar investigation or anogenital investigation of the familiar stimulus animal (Fig. 4E&F). During the first 1 min of the test, social discrimination (novel anogenital investigation time / novel + familiar anogenital investigation time x 100) was significantly decreased in LPS-treated females, as indicated by a trend in whole body social discrimination (F _(1, 33)_ = 3.983, p=0.05, Fig. 4G) primarily driven by a significant decrease in anogenital investigation (F _(1, 33)_ = 5.665, p<0.05, Fig. 4H). Only the saline-treated Cre-females showed a significant preference for the novel stimulus within the first one minute of introduction, as measured against 50% (chance; p<0.05; Fig. 4G), indicating a social ‘memory’ for the previously encountered familiar conspecific. Finally, no significant differences were observed in rearing, digging, or grooming behaviors (Table 4).

**Table 4.**
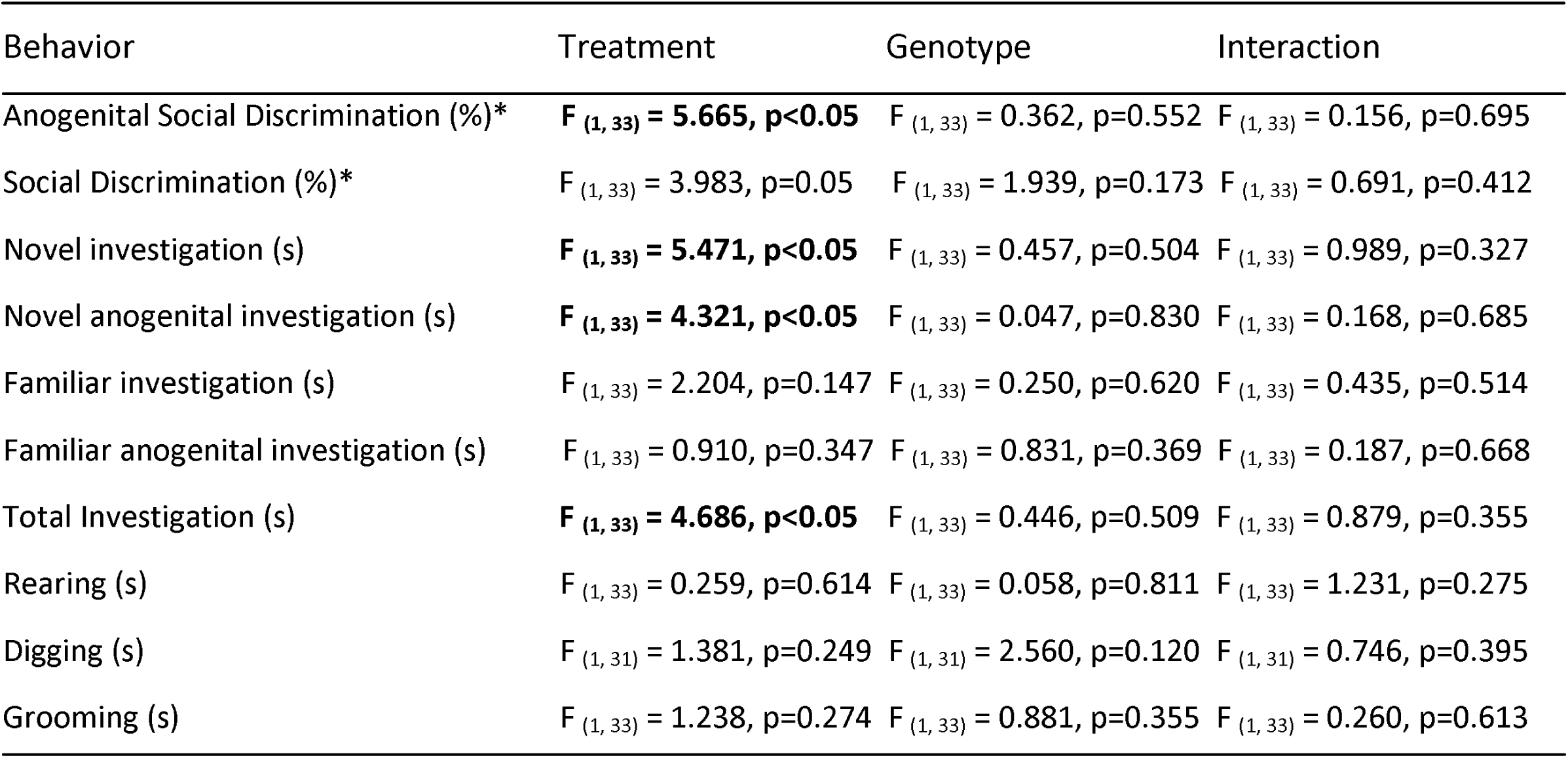
2-way ANOVA results of social discrimination testing in females. Bolded statistics indicate significant differences, * based on first 1 min. of the test, s:seconds.

**Figure 4.**
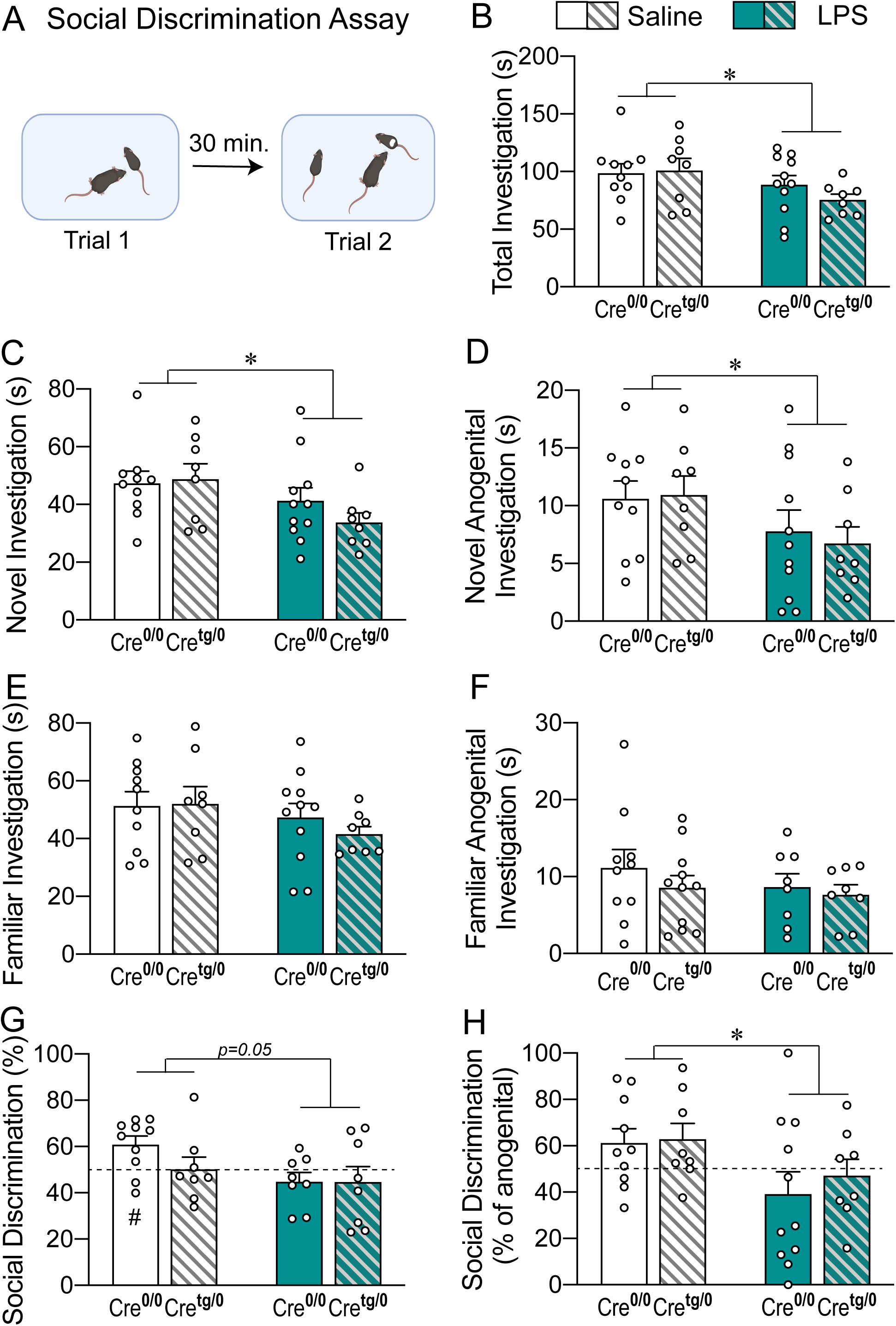
PND4 LPS challenge reduces social discrimination in adult females, independent of microglial-MyD88 signaling. **A)** Social Discrimination Assay in which the subject female is exposed to a novel female juvenile for 5 min (Trial 1). Following a 30 min interval, the subject is again exposed to the same novel female (now ‘familiar’) as well as a new novel juvenile female (Trial 2). Results shown are for social discrimination (Trial 2). LPS-treated females spend less time in total social investigation (**B**), novel investigation (**C**), and novel anogenital investigation (**D**) as compared to saline-treated females. No significant differences were found in familiar investigation (**E**) or familiar anogenital investigation (**F**). During the first minute of the test, only Cre^0/0^ Saline-treated females showed a significant preference for the novel stimulus (G) and social discrimination was reduced by neonatal LPS (**H**). *main effects of 2-way ANOVA (treatment x genotype), p<0.05. ^#^single sample t-test against 50% (chance), p<0.05.

**Figure 5.**
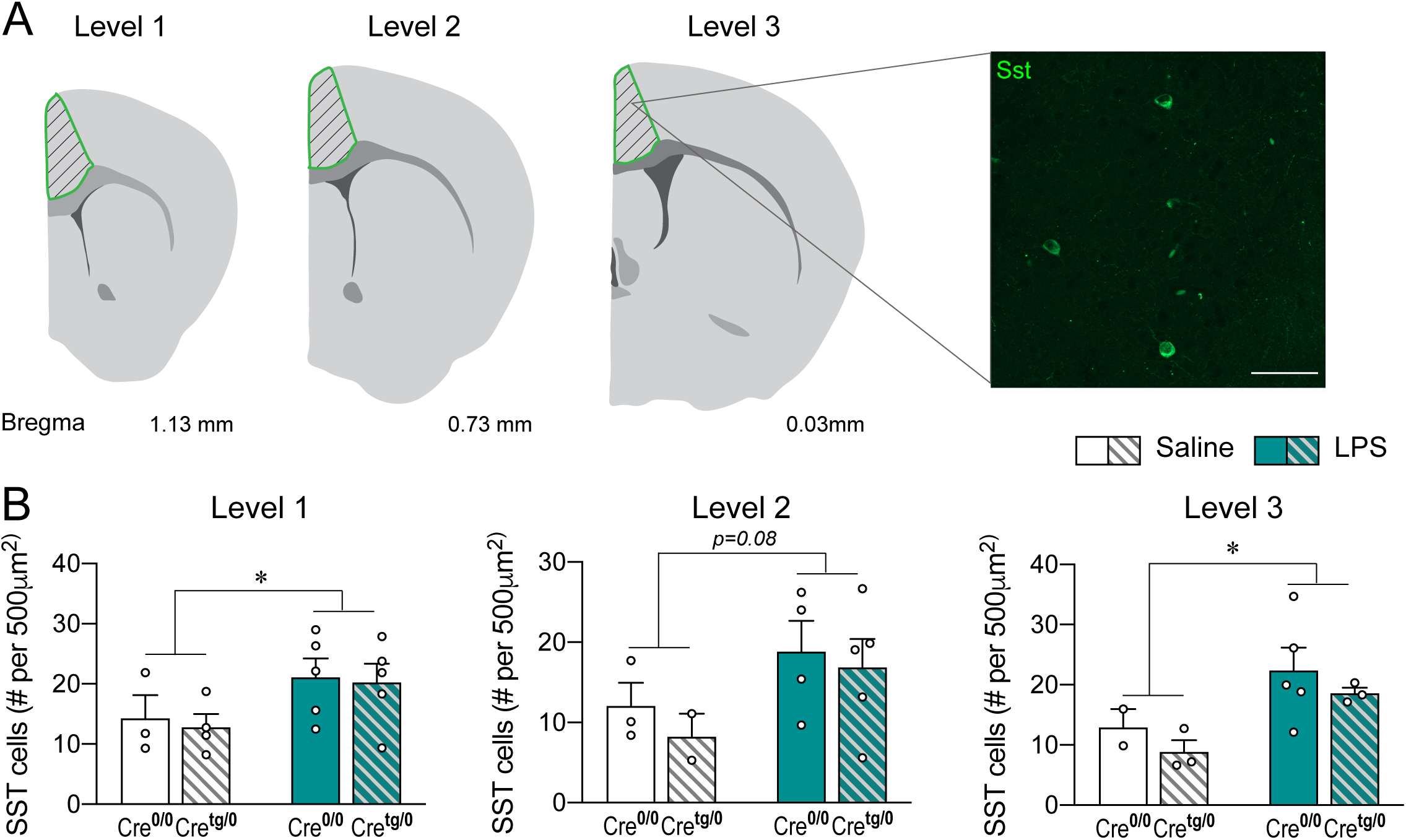
PND4 LPS challenge increases SST cell number in the ACC in females, independent of microglial-MyD88 signaling. **A**) Atlas image representations depicting the Bregma distances corresponding to each Level of SST analysis in the ACC, as well as representative image of immunohistochemical staining for SST. **B**) Neonatal LPS increases SST cell number in Levels 1 & 3, with a trend towards a significant effect in Level 2 (p=0.08). *main effects of 2-way ANOVA (treatment x genotype), p<0.05.

### 3.4 PND4 LPS challenge increases SST cell number in the ACC in females, independent of microglial-MyD88 signaling

Somatostatin cell number was significantly increased in the ACC of LPS-treated females as compared to saline-treated females, regardless of genotype (see Table 5 for complete statistics). Cell number was assessed at 3 different rostro-caudal levels (Bregma distances) throughout the ACC (Fig. 5A). Main effects of treatment reached statistical significance at Levels 1 and 3 (Level 1: F _(1, 13)_ = 4.999, p<0.05, Level 3: F _(1, 9)_ = 7.226, p<0.05; Fig. 5B) with a trend towards a significant increase at Level 2 (F _(1,_ 10) = 3.760, p=0.08; Fig. 5B), suggesting that this increase in SST cell number is widespread within the ACC. No significant effects of genotype were observed at any level.

**Table 5.** 2-way ANOVA results for Somatostatin cell number in the ACC. Bolded statistics indicate significant effects.

| Level | Treatment | Genotype | Interaction |
| --- | --- | --- | --- |
| Level 1 | **F<sub>(1, 13)</sub> = 4.999, p<0.05** | F<sub>(1, 13)</sub> = 0.140, p=0.715 | F<sub>(1, 13)</sub> = 0.010, p=0.922 |
| Level 2 | F<sub>(1, 10)</sub> = 3.760, p=0.08 | F<sub>(1, 10)</sub> = 0.534, p=0.482 | F<sub>(1, 10)</sub> = 0.057, p=0.817 |
| Level 3 | **F<sub>(1, 9)</sub> = 7.226, p<0.05** | F<sub>(1, 9)</sub> = 1.203, p=0.301 | F<sub>(1, 9)</sub> = 0.002, p=0.968 |

## 3. Discussion

In contrast to our initial hypotheses, our results show that a neonatal LPS challenge reduced sociability in adult female mice, while no sociability deficits were observed in adult males. LPS-treated females also displayed reduced social interaction and social memory in a social discrimination task, as well as an increase in SST cell number in the ACC. Finally, MyD88 knockdown significantly decreased TNFα and IL-1β mRNA in microglia following an LPS challenge, but did not prevent LPS-induced changes in either social behavior or SST cell number. Cumulatively, these results suggest that neonatal endotoxin exposure has long-lasting consequences for social behavior in females, independent of microglial-MyD88 signaling.

Our results demonstrate that neonatal LPS disrupts social behavior in adult females across multiple behavioral assays. To the best of our knowledge, we are the first to report decreases in sociability and social discrimination memory in adult females following neonatal endotoxin exposure. However, these findings are in line with previous studies showing that adult female social behavior is altered by immune activation. For example, neonatal LPS decreases social interaction time in female rats tested during adolescence (Doenni et al., 2016; MacRae et al., 2015). In Siberian hamsters, PND3 LPS-treated females investigate an opposite-sex conspecific more and spend more time in attack bouts than saline-treated females (Sylvia & Demas, 2017). LPS administration at later developmental timepoints also influences female social behaviors. In female CD1 mice, pubertal LPS decreases sexual receptivity in adulthood (Ismail et al., 2011) and while estradiol normally enhances social memory in a social discrimination task, this enhancement is prevented by pubertal LPS (Ismail & Blaustein, 2013). Similarly, acute LPS challenge in adult females has been shown to reduce nest building, pup retrieval, and maternal aggression in mice (Weil et al., 2006; Aubert et al., 1997) and facilitate partner preference in prairie voles (Bilbo et al., 1999). Interestingly, perinatal immune challenge also influences female reproductive physiology. In rats, neonatal LPS accelerated the onset of puberty, as evidenced by an earlier age of vaginal opening and estrous cycling onset (Sominsky et al., 2012). Corticosterone levels were also elevated in neonatal LPS-treated females across developmental timepoints (Sominsky et al., 2012). LPS-treated female Siberian hamsters have abnormal estrous cycles and smaller ovaries relative to saline controls, while male testes mass is not affected (Sylvia & Demas, 2017).

The sex-specificity of neonatal LPS effects on social behavior may also depend on a variety of factors, including drug dose, age at drug administration, and species. For instance, in mice treated with a high dose of LPS (10mg/kg) at PND9, Carlezon et al., 2019 observed sociability deficits in males only. Using a lower dose (50µg/kg), Custodio et al., 2018 observed no effect of neonatal LPS on social behavior in either male or female mice. In rats, 50-100 µg/kg of LPS at either PND3-5 or PND14 decreases sociability in both males and females during adolescence (MacRae et al., 2015; Doenni et al., 2016). When taken into consideration along with our current results, these findings may suggest that male mice display social behavior deficits following much higher doses of LPS (greater than the 330µg/kg that we used in this study), while female mice display social deficits at doses that more closely approximate those that have been shown to induce behavioral changes in female rats.

In addition to decreasing social behavior, we found that neonatal LPS increased SST cell number in the ACC in adult females. This finding is highly intriguing given that SST interneurons have recently been shown to play a role in the modulation of social behavior (Perez et al., 2019; Sun et al., 2020; Nakajimi et al., 2014). Furthermore, while studies in male mice have shown that lower SST interneuron number is associated with decreased social behavior (Bonini et al., 2016; Chen et al., 2019), here we observe the presence of *higher* SST interneuron number and lower social behavior in female mice. This could suggest multiple interesting possibilities. First, it might be the case that the regulation of social behavior by SST interneurons is sex-specific. Further work is needed to fully characterize the impact of neonatal LPS on SST interneurons in both males and females. Second, because SST interneurons in the ACC tightly regulate the activity of surrounding pyramidal neurons (Kapfer et al., 2007; Tan et al., 2008; Cardin, 2018), it is possible that either increases or decreases in their number or activity could inhibit the display of appropriate social behavior. Finally, parvalbumin (PV) interneurons in the ACC have also been implicated in the regulation of social behavior (Bicks et al., 2020; Deng et al., 2019; Selimbeyoglu et al., 2017), play a critical role in the modulation of overall neural activity in the ACC (Cardin, 2018; Hu et al., 2014), and are impacted by early life experience (Holland et al., 2014; Goodwill et al., 2018; Gildawie et al., 2020). Therefore, the expression of social behavior may depend on relative or coordinated activity of these two cell types. While answering these questions is beyond the scope of the current study, they suggest interesting avenues for future inquiry.

Based on the fact that microglia respond to immune challenges early in life (Bilbo & Schwarz, 2012; Williamson et al., 2011) and have the capacity to organize social circuits in the brain (Nelson et al., 2017; Kopec et al., 2018; VanRyzin et al., 2019; Ikezu et al., 2020), we hypothesized that removal of microglial-MyD88 signaling would prevent the LPS-induced changes in behavior and SST cell number. However, in contrast to this initial hypothesis, we observed no effect of MyD88 knockdown on any of our outcome measures. Yet, given that the behavioral effects that we observed were female-specific, this may not be so surprising. Much of the evidence suggesting a key role for microglia in these processes has been male-specific. For instance, Kopec et al. (2018) found that while social play behavior changes over the course of adolescence in both males and females, microglial sculpting of the underlying neural circuitry is specific to males. Indeed, the mechanism by which the developmental circuitry change occurs in females to modify social play behavior remains unclear. Thus, it may be the case that microglia are less critical to the modulation of female behaviors. Another possibility is that the LPS-induced behavior changes are mediated by the intact MyD88-independent pathway downstream of TLR activation in microglia. This seems unlikely, however, given that INFβ was significantly increased following LPS in males, but not in females. A final, intriguing possibility is that these social behavior deficits are driven by peripheral immune signaling. In line with this idea, peripherally administered LPS does not enter the brain directly, but leads to a robust induction of cytokines such as TNFα which then induce neuroinflammation (Qin et al., 2007). While the majority of TNFα receptors within the brain are located on microglia, they are also expressed by other cell types such as astrocytes, endothelial cells, and oligodendrocyte progenitor cells (Brainrnaseq.org). LPS also has the capacity to impact T cell differentiation (Koch et al., 2007; McAleer & Vella, 2008) and T cells have, in turn, been shown to mediate behaviors such as anxiety-like behavior in mice by acting on oligodendrocytes (Fan et al., 2019). Similarly, interferon gamma has been shown to mediate social behavior by increasing inhibitory tone in cortical neurons (Filiano et al, 2016).

Collectively, our results demonstrate that neonatal exposure to LPS leads to deficits in both sociability and social discrimination in adult females, as well as increased SST interneuron number in the ACC. Moreover, they suggest that these effects are not mediated by MyD88-dependent TLR signaling in microglia. We hope that these findings may serve as a springboard for future studies aimed at better understanding the impact of early life immune challenge on female neuroimmune function and behavior.

## Acknowledgements

We would like to thank Drs. Evan Bordt and Alexis Ceasrine for their careful and critical reading of the manuscript.

## Funding Sources

This work was supported by R01MH101183 to S.D.B and F32ES029912 to C.J.S. and by the Robert and Donna E. Landreth Family Fund.

## Notes

### Competing Interest Statement

The authors have declared no competing interest.

